# Advancing clinical outcome predictions via incorporating pharmacokinetic simulations into *in vitro* testing - a colorectal cancer example

**DOI:** 10.1101/2025.07.29.663668

**Authors:** Andrey A. Poloznikov, Ben R. Britt, Sergey Nikulin, Sergey Rodin, Karl-Henrik Grinnemo, Martin Woywod, Jasmin Farouq

## Abstract

The development of *in vitro* assays that can predict clinical outcomes is highly desirable for drug development and personalized medicine. However, conventional *in vitro* methods often fail to replicate physiological drug pharmacokinetics, posing a challenge to their clinical translation. To address this issue, we adjusted incubation times and concentrations of standard-of-care drugs in the *in vitro* chemosensitivity assay to reflect those encountered by colorectal cancer patients. Then, for first time, we mimicked the relevant drug exposure of mFOLFOX-6, CapOx and FOLFIRI protocols to predict clinical outcomes. Our pharmacokinetic-based testing on primary colorectal cancer cells accurately predicted responders and non-responders among a cohort of patients (N=6).

Classical testing methods such as IC_50_ and GI_50_ did not reveal any clinically meaningful results. Furthermore, we demonstrated that even subtle changes in drug incubation times could lead to significant variations in the classification of cells as sensitive and resistant, which is not related to mechanisms of action according to categorical clustering. Finally, our pharmacokinetic-based test results were consistent with the historical clinical data on similarities of mFOLFOX-6 and CapOx schemes.

Our results contribute to the growing body of evidence that pharmacokinetic-based *in vitro* testing could bridge the gap between laboratory research and clinical practice. Integration of pharmacokinetic dynamics into *in vitro* tests could have a significant potential in enhancing drug development and refining personalized treatment strategies.

## Introduction

The development of an *in vitro* assay to predict clinical outcomes is one of the main research directions in the fields of drug development and personalized medicine. In the early stages of drug development, drug candidates’ efficacy and toxicity are tested in standardized cellular assays, enabling higher throughput. IC_50_ and EC_50_ (1), which are derived from dose-response curves in these assays, are essential metrics to study the structure-activity relationship of tested compounds and identify molecular characteristics correlated with drug sensitivity (2–4). With the development of patient-derived organoids technology (5), IC_50_ values were also used as a biomarker to predict response to treatment in patients (6). However, as recently analyzed in (7), developing individualised tumor response tests using patient-derived organoids (PDOs) remains a challenging task.

The clinical validity of PDO-based tests can vary depending on the characteristics of the tumor and the treatments being tested. Despite promising results in colorectal and pancreatic cancer, data from patients with gastric and esophageal cancer do not convincingly show an association between PDO drug test results and clinical response. Furthermore, the results are predictive for patients who receive treatments based on irinotecan and gemcitabine, while there are conflicting results for oxaliplatin-based treatments (8–11). We hypothesized that these limitations and inconsistencies can be attributed to the differences in response metrics and testing conditions used in the studies.

Although IC_50_ is the most widely used metric to analyze dose-response curves, it is not the most reliable of the metrics to quantify drug sensitivity (12). Better performance was shown for the fitted or trapezoidal area under the dose-response curve (AUC), and this parameter was used in ca. 30% of the studies mentioned in the review by Wensink et al. Wensink et al. (7). An alternative drug response metric, that is insensitive to division number, is growth rate inhibition (13).

Since IC_50_ values of cell cycle-dependent cancer drugs, including DNA synthesis inhibitors (i.e. 5-fluorouracil, gemcitabine, platinum-containing drugs) and microtubule inhibitors (i.e. taxanes, vinblastine) are highly sensitive to the number of divisions that occur over the course of a response assay, GR metrics were shown to be superior to assess the effects of these compounds on dividing cells (14). Other metrics used were decrease in viability, organoid size, and metabolic score (7). Tan et al. Tan et al. (15) presented a pan-matrix viability score that yielded promising results in colorectal cancer for combination therapy. With a plethora of metrics mentioned above, the optimal one still needs to be defined.

Another dimension of optimization that is needed to achieve clinically relevant results is the experimental setup. Conventional testing approaches poorly replicate the physiological pharmacokinetics of a drug seen *in vivo* that hamper the translation of *in vitro* parameters into clinical estimations.

One of the aspects is the exposure time. The absence of guidelines for selecting this parameter can lead to high variability in the *in vitro* data obtained (16). Among the parameters that should be considered when estimating assay time are the cell doubling time and stability of a drug in solution (17). The assay time should be long enough to allow tracking the difference in cell viability between control and experimental samples that increases with a number of cell divisions that occurred starting from the zero time point. The example of oxaliplatin clearly illustrates the importance of measuring the stability of compounds in the culture medium used. As demonstrated by Jerremalm et al. Jerremalm et al. (18), oxaliplatin undergoes rapid degradation in solutions containing chloride ions such as plasma and culture medium. Therefore, a long incubation time could lead to skewed results (9–11). On the other hand, the exposure time of drugs varies widely in patients and is strongly influenced by half-life time (19) with median values of 0.48 days for small molecules and 4.82 days for biologicals (Figure 1). In organoids the most common incubation time was 6 days regardless of individual characteristics of the drugs that led to overtreatment of the cells with small molecules (Figure 2). Although a longer incubation time results in a shift of a dose-response curve to the lower concentration region, the degree of such shift should be determined in each particular case (20).

**Fig. 1.**
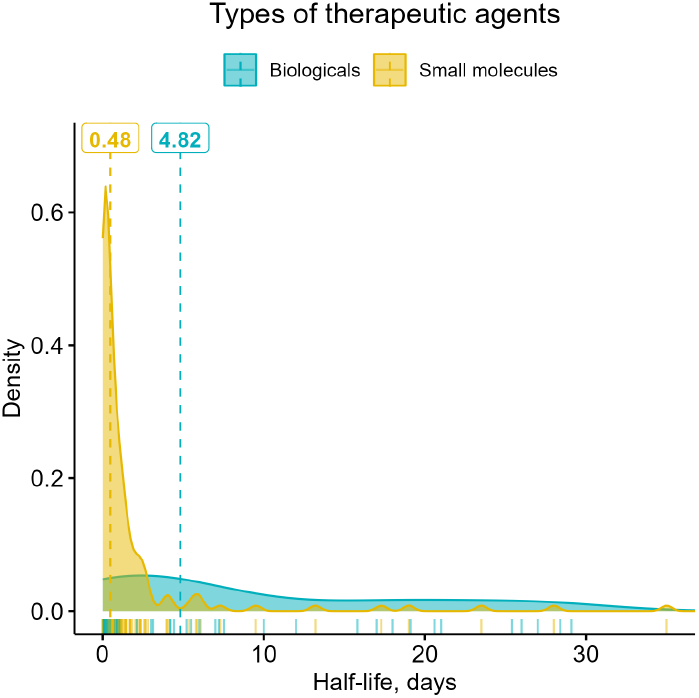
Distribution chart of half-life times for cancer drugs (19).

**Fig. 2.**
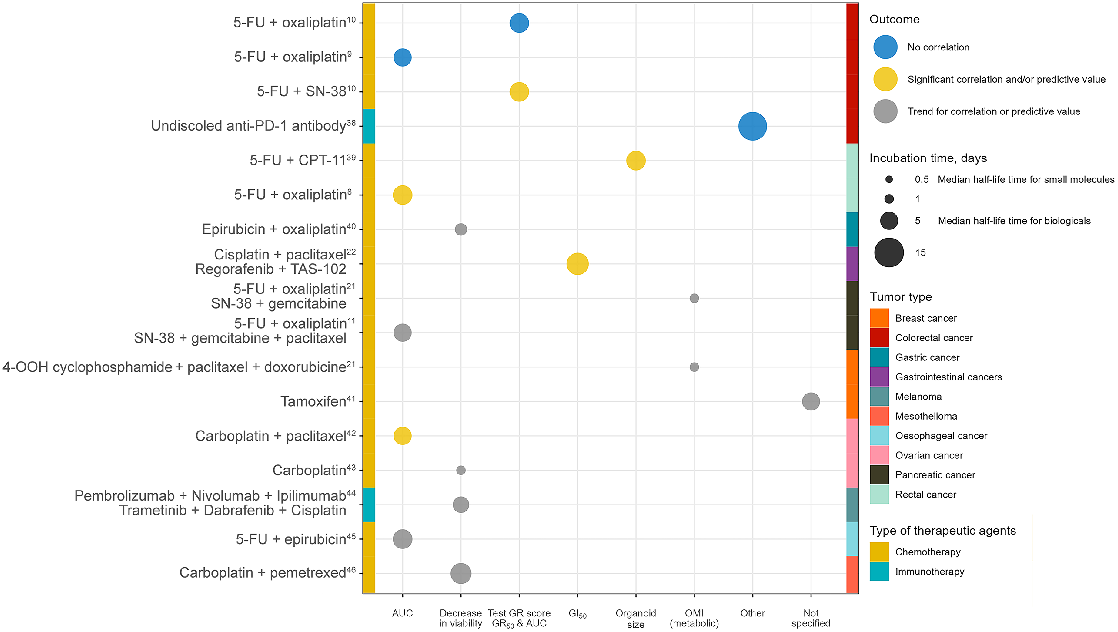
Assay parameters and outcomes in PDO studies reported (8–11, 21, 22, 38–46).

The second aspect of the experimental setup is concentration. In dose response assays cells are often exposed to drug concentrations that exceed clinically achievable levels. Only five out of 18 studies reviewed referenced maximum plasma concentrations (C_max_) values (9, 10, 15, 21, 22). At the same time, the clinical validity of the IC_50_ and EC_50_ values that exceed C_max_ without any further normalization is questionable (23, 24).

In oncology, most treatment schemes include drug combinations that are administered according to a certain schedule (25). Although in published PDO studies the drugs are usually incubated together, if the drugs in a combination exhibit synergism, antagonism, or additive effect, the overall effect could be schedule dependent (26, 27). For example, the effects of the ternary combination of irinotecan, 5-fluorouracil, and oxaliplatin, which is a standard of care treatment for colorectal cancer, could be antagonistic or synergistic in the different schedules tested (28). Therefore, recapitulation of the drug administration sequence is another aspect of the experimental setup that is worth considering.

In general, current *in vitro* methods represent a static ‘closed’ pharmacodynamic model that requires multidimensional optimization of the parameters to overcome the limitations mentioned above. Another approach to improve clinical validity is to simulate an ‘open’ *in vivo* system. This requires that receptor occupancy, target turnover, and target-drug interactions dynamics are included in the model (29, 30). One of the first steps in this direction could be to mimic the pharmacokinetic (PK) profile of a drug or drug combination according to the treatment scheme. With the availability of the PK data for the marketed drugs, this could bring more physiological relevance to the *in vitro* experiments. Previous studies have demonstrated that imitation of bolus and continuous exposure not only lead to different response rates but also have an impact on the development of drug resistance (31–34). Finally, incorporation of PK exposures into cellular assays was demonstrated to improve translation in xenograft models (35–37).

We aim to develop an *in vitro* assay that can more accurately predict how cancer patients will respond to standard treatments. In this paper, we are addressing thus the following research objectives:

1. **Address limitations of current *in vitro* methods:** We highlight the importance of replicating physiological conditions in *in vitro* assays by illustrating how the drug exposure time impacts the sensitivity status of cancer cells.
2. **Develop *dynamic dose* protocols:** We create *in vitro* protocols that mimic the pharmacokinetics of standard colorectal cancer treatments (specifically, the mFOLFOX-6, CapOx, and FOLFIRI schemes), aiming to replicate clinical exposure dynamics observed in patients.
3. **Compare *in vitro* results with clinical outcomes:** We validate the developed *in vitro* assays by comparing the results with responses of colorectal cancer patients, thereby underlining the relevance and applicability of our findings.

## Material and methods

Human colorectal cancer (CRC) cells were cultured in a complete RPMI 1640 (COLO 205) or DMEM high glucose (HT-29, HCT 116, SW480 and SW620) cell culture medium (Gibco, USA) supplemented with 10% vol. fetal bovine serum, 1% vol. GlutaMax (Gibco, USA) and 1% vol. antibiotic-antimycotic solution (Gibco, USA). Cells were incubated in a cell culture incubator (37°C, 5% CO_2_). Subculture was carried out every 2-3 days using TrypLE dissociation reagent (Gibco, USA). Primary colorectal cancer cells were derived previously upon written informed consent of the respective patients (47, 48) and cultured in advanced DMEM/F12 (Gibco, United States) supplemented with 1% vol. (Gibco, USA), 1% vol. antibiotic-antimycotic solution (Gibco, USA), 1% vol. HEPES (Gibco, USA), 2% vol. B27 (Gibco, USA), 1.25 mM N-acetylcysteine (Sigma, USA), 10 mM nicotinamide (Sigma, USA), 250 ng/ml R-spondin 1 (PeproTech, USA), 100 ng/ml noggin (PeproTech, USA), 50 ng/ml human EGF (Gibco, USA), 10 nM gastrin I (Sigma, USA), 500 nM A83-01 (STEMCELL Technologies, Canada), 1 *µ*M SB202190 (Tocris Bioscience, UK), 10 nM prostaglandine E2 (Sigma, USA), and 5 *µ*M Y-27632 (STEMCELL Technologies, Canada) in a Matrigel GFR basement membrane matrix domes (Corning, USA). Cell culture medium was replaced every 48h. Subculture was carried out every 2 weeks using TrypLE dissociation reagent (Gibco, USA). Cells were counted after trypan blue staining (Gibco, USA) using an EVE automated cell counter (NanoEntec, Korea) according to the manufacturer’s protocol.

### Drug test

HT-29, HCT 116, COLO 205, SW480 and SW620 cells were seeded in 96-well plates (TPP, Switzerland) in 100 ul of complete cell culture medium (5000 per well). Primary cultures were diluted in 10 *µ*l Matrigel GFR (Corning, USA) and seeded in 96-well plates (TPP, Switzerland) (50 spheroids per well). After solidification of the gel, 100 ul of complete cell culture medium was added to each well. After 24h, cell culture medium was replaced with medium containing oxaliplatin, 5-fluorouracil (5-FU) and SN-38 (Tocris Bioscience, UK) respectively. To determine IC_50_ and GI_50_ values, cells were exposed to serial dilutions covering physiological concentrations of 5-FU (5-FU C_max_ in patients = 341 *µ*M (49); 5-FU range *in vitro* = 0.26 to 800 *µ*M), oxaliplatin (oxaliplatin C_max_ in patients = 10 *µ*M (50); oxaliplatin range *in vitro* = 0.19 to 200 *µ*M) and SN-38 (SN-38 C_max_ in patients = 86 nM (51); SN-38 range *in vitro* = 0.32 to 1000 nM). 5-FU and SN-38 were prepared in DMSO, while oxaliplatin stock solution was prepared in water (52). Treatments were compared with vehicle controls that contained the same amount of DMSO and water. Cells were incubated for 72 hours in a cell culture incubator (37°C, 5% CO_2_) and the medium was replaced with a complete fresh cell culture medium. To evaluate the sensitivity of cancer cells to CRC treatment schemes, we generated unique PK-based *in vitro* testing protocols, to which we will henceforth refer as *dynamic dose* testing protocols, as they mimic the sequence, duration of exposure, and drug concentrations of the treatments. To do so, we referenced the hematology and oncology chemotherapy manual (25) to identify the schedule and timing of drug administrations. Additionally, we analyzed publicly available data on the pharmacokinetics of the drugs included in relevant treatment schemes. For example, the mFOLFOX-6 *dynmaic dose* protocol includes a 2-hour incubation with 3.37 *µ*M of oxaliplatin, a 30-minute incubation with 94.5 *µ*M of 5-FU, and a 46-hour incubation with 3.4 *µ*M of 5-FU. Similarly, the CapOx *dynmaic dose* protocol includes a 2-hour incubation with 5.15 *µ*M of oxaliplatin, followed by two daily 3-hour incubations with 10 *µ*M of 5-FU for three days. And finally, FOLFIRI *dynmaic dose* protocol includes a 1.5-hour incubation with 50.7 nM of SN-38, a 30-minute incubation with a mixture of 94.5 *µ*M of 5-FU and 50.7 nM of SN-38, a 2.5-hour incubation with a mixture of 3.4 *µ*M of 5-FU and 50.7 nM of SN-38, an 18.5 hour incubation with a mixture of 3.4 *µ*M of 5-FU and 21.1 nM of SN-38, and a 26-hour incubation with 3.4 *µ*M of 5-FU. After the last incubation, the medium was replaced with fresh complete cell culture medium and the cells were incubated in a cell culture incubator (37°C, 5% CO_2_) for a total of 72 hours from the start of the assay. A detailed description of these protocols is provided in the results section. Cell viability was measured with the CellTiter 96 aqueous solution cell proliferation assay kit (Promega, USA) according to the manufacturer’s instructions (40). The cell growth rate was calculated as described previously (13).

### Data analysis

For the analysis of the effect of exposure time on the classification of cells as sensitive or resistant to individual drugs, we used previously published data (20) available online at: https://brb.nci.nih.gov/ETvsCT/. The parameters ‘EC_50__Estimate’, ‘Slope_Estimate’, ‘LowerLimit_ Estimate’ and ‘UpperLimit_Estimate’ were utilized to determine the area under dose-response curves (AUC) for all drugs tested and normalized to the control. We set the median of the normalized AUC values distribution as threshold. Cells were classified as ‘sensitive’ or ‘resistant’ to a particular drug under different incubation times based on whether their normalized AUC values were above or below this cutoff. To assess the overall impact of exposure time on cell classification, we normalized the number of cells that retained their sensitivity or resistance to a drug to the total number of cell-drug pairs tested across all incubation times. The calculations were performed using R language (53) in the RStudio environment (54) and dplyr (55), tidyr (56), stringr (55) packages. The graphs were plotted in GraphPad Prism software (GraphPad Prism Software, USA) and ggplot2 (57), ggsci (58), ggh4x (59), patchwork (60) R packages. For the categorical clustering we used the Python module KModes v0.12.2 (61, 62). Essentially, the algorithm is initialized with *k* random modes, representing the clusters characteristic cell response. Then the drugs are assigned to the cluster with the corresponding mode which shares the most amount of same cell responses. Each mode is then updated to the most frequent cell response of all drugs in the corresponding cluster. These two steps of assigning and updating are then repeated until convergence is reached, i.e. the mode of each cluster represents the most frequent cell response of all drugs in that cluster. We considered as final the converged clustering result with the least amount of dissimilarities between modes and corresponding drugs out of 1000 random initializations.

## Results and discussion

### Exposure time - sensitivity status relationship

Several factors have an impact on the response of cells to drugs, and the duration of exposure is critical. In the Evans et al. Evans et al. (20) study 320 compounds were screened in the NCI60 cell line panel using 2, 3, 7, and 11 days exposure times respectively. Although a positive linear relationship was observed between IC_50_ values and exposure, it was not clear whether the individual sensitivity status of the cells remains similar at different incubation times. Understanding this possible dependency of incubation time and drug sensitivity could play an important role in selection of assay parameters and further clinical decision making, e.g. personalized treatment recommendations after PDO testing. First, we calculated normalized to control AUC values for the 16225 concentration-response curves present in the dataset. Then we set the median of the normalized AUC values distribution as a threshold for each drug. Cells were classified as ‘sensitive’ or ‘resistant’ to a particular drug in a 2-day incubation assay. Surprisingly, we observed that even small changes in incubation time (3 days vs. 2 days) led to significant changes in the order of cell line sensitivities (Figure 3A). As clearly seen in the 5-FU example, the cells that were labeled as ‘sensitive’ in a 2-day incubation assay were distributed across all ‘Normalized AUC’ axis range in a 3-day incubation assay without any clear new cutoff value. Similar observations were found for all drugs (data not shown).

**Fig. 3.**
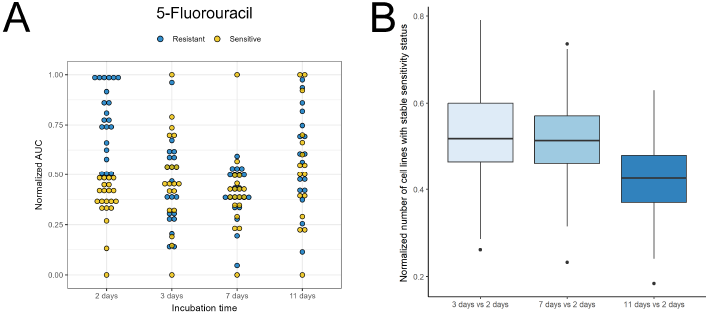
(A) Distribution of ‘sensitive’ and ‘resistant’ cell lines to 5-FU based on 2-day incubation data after 3, 7, and 11 days of incubation. (B) Normalized number of cell lines that maintain their 2-day incubation sensitivity status after 3, 7, and 11 days of incubation.

To assess the overall impact of exposure time on cell classification, we normalized the number of cells that retained their position relative to the median values to the total number of cell drug pairs for each incubation time (Figure 3B). The median value of cells that preserved the sensitivity status compared to 2-day incubation after 3 and 7-day incubation was roughly 0.5. More profound changes in the sensitivity status were found for cells incubated for 2 days and 11 days. However, this discrepancy could be attributed to the differences in culture methods, as the data for the 11-day incubation was acquired in spheroid culture, unlike the monolayer cultures for the other incubation times.

One of the possible explanations for the change in sensitivity status could be the mechanism of the action of drugs. For compounds that inhibit growth rate or exhibit cytotoxic effect in a cell cycle-dependent manner, the IC_50_ or EC_50_ values are highly sensitive to the number of divisions that occur during exposure time (13). Such compounds include selective S-phase selective agents, DNA-damaging agents, DNA methyltransferase inhibitors, DNA polymerase inhibitors and intracellular serine-threonine kinase inhibitors. On the other hand, histone deacetylase inhibitors, receptor tyrosine kinase inhibitors, EZH2 inhibitors, gamma secretase inhibitors, and SMO inhibitors had no dependence on exposure time in manifesting growth inhibition or were not effective on cell viability even after 11-day exposure (20).

To assess whether the mechanisms of action affect the preservation of the sensitivity state and to identify the ‘exposure-agnostic’ drugs, we compared the number of cells with consistent sensitivity states between different drug types (Figure 4, Supplementary data file 1).

**Fig. 4.**
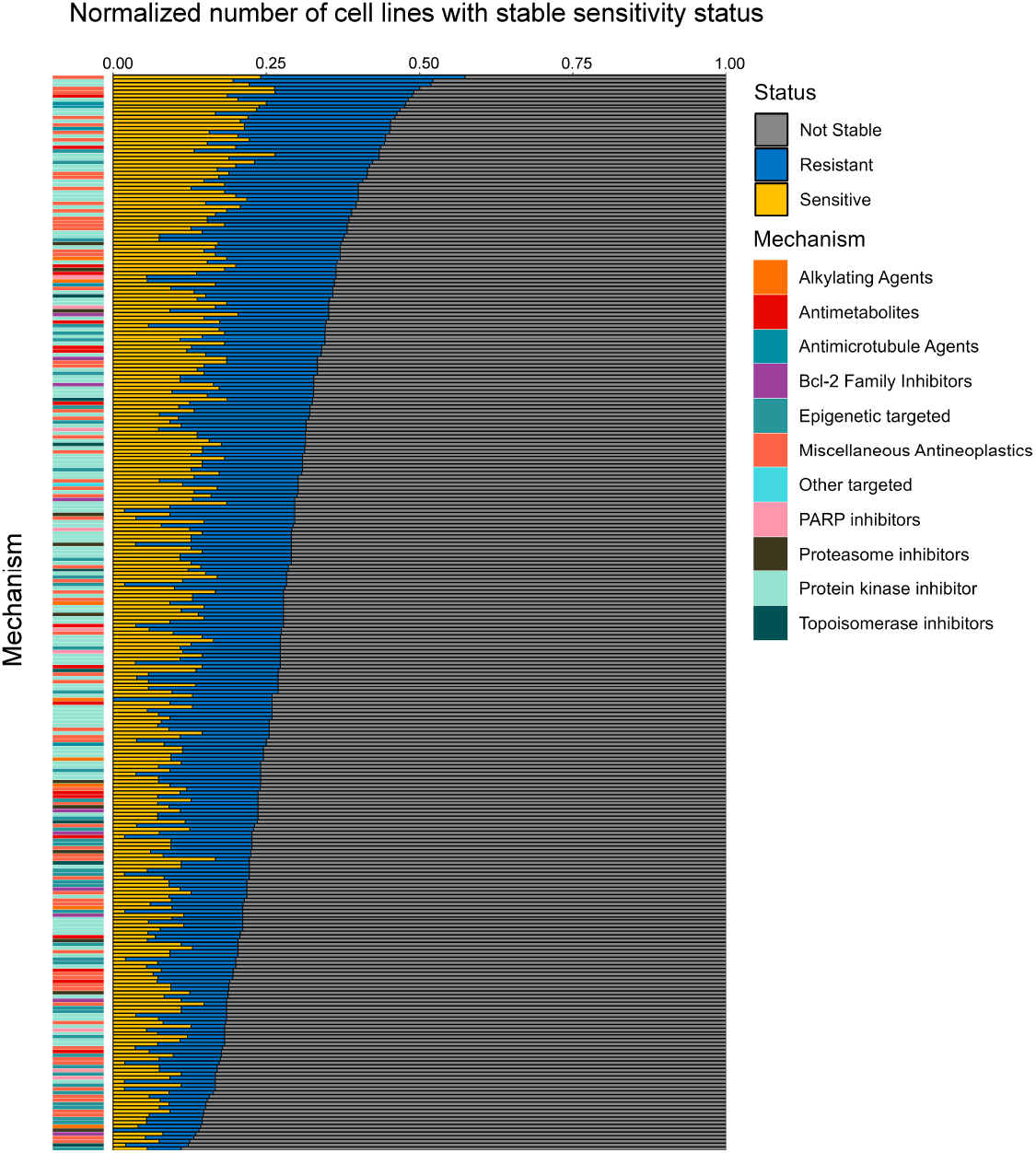
Normalized number of cell lines that had a consistent sensitivity status at all incubation times with a drug.

Although the range of cells with stable sensitivity varies between 10% and 55% for different drugs, an obvious explanation for the sensitivity status of the cells could not be found based on the mechanism of action of the drugs. Therefore, we used *k*-mode clustering in attempt to derive such drug clusters. The cell lines were labeled according to the their response status to a drug across different incubation times (‘sensitive’, ‘resistant’, ‘unstable’).

As *k*-mode clustering relies on a predefined number of clusters, we tried from *k* = 4 to 11, representing the number of mechanisms of action into which the tested drugs can be grouped (Supplementary data file 2). To interpret the results, we carried out a *k*-mode analysis and compared the derived clusters with the categorisation according to the known mechanism of action of drugs as visualised in figure 5 for *k* = 4.

**Fig. 5.**
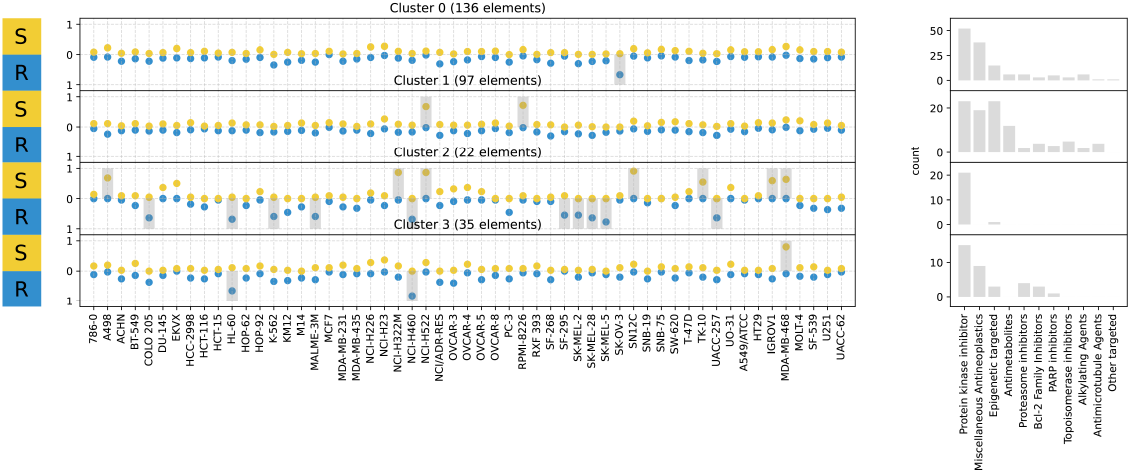
Mode Analysis of *k*-mode clustering of cell responses (left) and comparison with know mechanism of action of the drugs (right). The modes of each cluster are indicated by the up or down facing grey bars for sensitive and resistant cell responses (missing for unstable cells). The yellow and blue circles indicate the relative frequency of sensitive and resistant behaviour of each cell lines.

The only insight that might be drawn is that drugs in cluster 3 are almost all protein kinase inhibitors. However, the the drugs with such mechanism of action were also present in other clusters. In any case, the results did not allow to identify any ‘exposure-agnostic’ category of drugs.

In addition to the factors mentioned above, drug stability in the cell culture medium may also account for these results. As shown in the example of platinum complexes, the presence of DMSO and chlorine ions may have an impact on actual concentrations of drugs in the assay (18, 52). Furthermore, the presence of DMSO and serum can affect the response to compounds with limited aqueous solubility (30). Therefore, we hypothesized that considering individual drug characteristics and pharmacokinetics in *in vitro* tests, i.e. using the *dynamic dose* approach, could improve the clinical validity of the corresponding data. To validate this hypothesis, we generated *dynamic dose* testing protocols for CRC treatment schemes and assessed the viability of CRC cell lines and primary colorectal cancer cell cultures in static and dynamic exposure assays.

### Generation of *dynamic dose* protocols for the FOLFOX-6, CapOx and FOLFIRI schemes

To develop *dynamic dose* protocols, we referred to the clinical recommendations (25). The main chemotherapy treatment options for the CRC patient are platinum-containing (FOLFOX and CapOx) or irinotecan-based (FOLFIRI) schemes. The sequences and doses of the drug constituting these schemes are presented in Table 1. We analysed the available PK data to determine the relevant *in vitro* concentrations individually for each scheme.

**Table 1.** Clinical schedules for the FOLFOX-6, CapOx and FOLFIRI schemes.

| Day | Timing | Drug | Dose | Route | Infusion duration |
| --- | --- | --- | --- | --- | --- |
| mFOLFOX-6 |  |  |  |  |  |
| 1 | 0 | Oxaliplatin | 85 mg/m<sup>2</sup> | i.v. | 2h |
| 1 | +2h | Calciumfolinat/Leukovorin® | 400 mg/m<sup>2</sup> | i.v. | 30min |
| 1 | +2h 30min | Fluorouracil (5-FU) | 400 mg/m<sup>2</sup> | i.v. | B |
| 1 | +2h 35min | Fluorouracil (5-FU) | 2400 mg/m<sup>2</sup> | i.v. | 46h |
| CapOx |  |  |  |  |  |
| 1-14 | 1-0-0-0 | Capecitabin | 1000 mg/m<sup>2</sup> | p.o. | - |
| 1-14 | 0-0-1-0 | Capecitabin | 1000 mg/m<sup>2</sup> | p.o. | - |
| 1 | 0 | Oxaliplatin | 130 mg/m<sup>2</sup> | i.v. | 2h |
| FOLFIRI |  |  |  |  |  |
| 1 | 0 | Irinotecan | 180 mg/m<sup>2</sup> | i.v. | 1h 30min |
| 1 | +1h 30min | Calciumfolinat/Leukovorin® | 400 mg/m<sup>2</sup> | i.v. | 30min |
| 1 | +2h | Fluorouracil (5-FU) | 400 mg/m<sup>2</sup> | i.v. | B |
| 1 | +2h 10min | Fluorouracil (5-FU) | 2400 mg/m<sup>2</sup> | i.v. | 48h |

i.v. – intravenous infusion; p.o. – oral administration; B – bolus infusion

### Dynamic dose mFOLFOX-6 protocol

#### Oxaliplatin infusion

Most pharmacokinetic studies on oxaliplatin primarily report data on platinum content rather than the drug itself, with measurements often taken in whole plasma, ultrafiltrate and erythrocytes (50, 63). However, it is important to note that oxaliplatin undergoes spontaneous metabolism, resulting in the platinum concentration not accurately reflecting the concentration of oxaliplatin. Additionally, the metabolites of oxaliplatin are either inactive or present at significantly lower concentrations than the parent compound, emphasizing the significance of considering the oxaliplatin content when determining *in vitro* concentrations (64). Studies conducted by Ehrsson et al. Ehrsson et al. (65), and Ehrsson and Wallin Ehrsson and Wallin (66), employed derivatization to measure oxaliplatin concentration, revealing an average maximum concentration of 1.44 ± 0.20 *µ*g/ml at a dose of 85 mg/m^2^, with the area under the PK curve (AUC^PK^) of 161 ± 23 *µ*g · min/mL. Consequently, for *in vitro* testing, a concentration of 3.37 *µ*M and an incubation time of 2 hours were selected. Notably, the incubation was performed in DPBS with 10 mM HEPES, as prior research demonstrated that after 2 hours in DPBS, around 80% of oxaliplatin remains in an intact state (18).

#### 5-FU bolus

The pharmacokinetics of 5-fluorouracil (5-FU) following bolus administration in doses of 300-600 mg/m^2^ have been extensively studied, showing peak concentrations in the millimolar range and rapid subsequent decline, with AUC^PK^ values ranging from 71 to 125 *µ*M · h (49, 67–69). For our *in vitro* experiment, we selected the AUC^PK^ data for the first 30 minutes after administration, as more than half of the AUC^PK^ values fall within this range (Collins et al., 1980). In this case, the AUC^PK^ has been identified to be 47.25 *µ*M · h (68). Thus, we chose a concentration of 94.5 *µ*M and an incubation time of 30 minutes to ensure a short exposure of high-concentration 5-FU to the cells.

#### 5-FU infusion

The *in vitro* concentration for simulating 5-FU infusion was determined based on the available data on mean 5-FU concentration and AUC^PK^. A mean AUC^PK^ of 20.35 *µ*M · h was identified for a 46-hour infusion, resulting in an *in vitro* concentration of 3.4 *µ*M and an incubation time of 46 hours (70). These findings align with 5-FU plasma concentration measurements (71).

### Dynamic dose CapOx protocol

#### Oxaliplatin infusion

Considering a linear increase in C_max_, the C_max_ for a dose of 130 mg/m^2^ is 2.2 *µ*g/ml (397.294 g/mol, 5.5 *µ*M). Additionally, the AUC^PK^ was determined to be 161 ± 23 *µ*g·min/mL for a dose of 85 mg/m^2^, which, assuming linearity, corresponds to 246 *µ*g·min/mL = 4.1 *µ*g·h/mL = 10.3 *µ*M·h. Therefore, for *in vitro* testing, a concentration of 5.15 *µ*M and an incubation time of 2 hours were selected. The incubation was conducted in DPBS with 10 mM HEPES, as it has been previously recommended for the mFOLFOX-6 protocol.

#### Capecitabine

Capecitabine (5’-deoxy-5-fluorouridine, 5’-DFUR) is a prodrug that undergoes conversion to 5-fluorouracil primarily in the liver. Subsequent metabolism leads to the formation of 5-fluoro-2’-deoxyuridine monophosphate (FdUMP) and 5-fluorouridine triphosphate (FUTP) in both normal and tumor cells. After a single dose of capecitabine at 1250 mg/m^2^, the AUC^PK^ for 5-FU reaches 4.4 *µ*M · h over a 6-hour period (72, 73). Approximately 90% of the AUC^PK^ is achieved within 3 hours. In the CapOx protocol, capecitabine is administered at a dose of 1000 mg/m^2^, resulting in a concentration of 1.2 *µ*M after a 3-hour exposure. However, when capecitabine is administered, 5-FU tends to accumulate in tissues. In the case of liver metastases, the tissue concentration of 5-fluorouracil can be nearly 10 times higher than the plasma concentration (73). Therefore, to simulate a single dose of capecitabine *in vitro*, a concentration of 10 *µ*M and an incubation time of 3 hours were selected. Considering the dosing schedule, this incubation was performed twice daily.

### Dynamic dose FOLFIRI protocol

#### Irinotecan infusion

Irinotecan is classified as a prodrug that undergoes liver metabolism to its active form, SN-38. Therefore, in our *in vitro* tests, we included SN-38. Based on a single-dose infusion of irinotecan at 180 mg/m^2^, the half-life of SN-38 was 11.02 hours (51). The calculated area under the PK curve was 269 ng · h/mL for SN-38. Considering the relatively long half-lives, we decided to simulate high and low concentration exposures through two separate incubations: 5.5 hours with 50.7 nM (SN-38), and 18.5 hours with 21.1 nM (SN-38). Furthermore, considering the subsequent administrations of 5-FU during treatment, we prepared double mixtures of two drugs to mimic their simultaneous presence in the bloodstream of patients.

#### 5-FU bolus

5-FU concentration of 94.5 *µ*M and an incubation time of 30 minutes as in the *in vitro* mFOLFOX-6 protocol.

#### 5-FU infusion

5-FU concentration of 3.4 *µ*M and an incubation time of 46 hours as in the *in vitro* mFOLFOX-6 protocol.

The resulting mFOLFOX-6, CapOx and FOLFIRI *dynamic dose* protocols are presented in Table 2. These protocols along with standard dose-response assays were used further to assess the sensitivity of colorectal cancer cells.

**Table 2.** *Dynamic dose* protocols for the FOLFOX-6, CapOx and FOLFIRI schemes.

| Day | Timing | Drug | Concentration | Incubation time |
| --- | --- | --- | --- | --- |
| *Dynamic dose mFOLFOX-6* |  |  |  |  |
| 1 | 0 | Oxaliplatin | 3.37 $\mu\text{M}$ | 2h |
| 1 | +2h 30min | Fluorouracil (5-FU) | 94.5 $\mu\text{M}$ | 30min |
| 1 | +3h | Fluorouracil (5-FU) | 3.4 $\mu\text{M}$ | 46h |
| *Dynamic dose CapOx* |  |  |  |  |
| 1 | 0 | Oxaliplatin | 5.15 $\mu\text{M}$ | 2h |
| 1-3 | +2h 30min | Fluorouracil (5-FU) | 10 $\mu\text{M}$ | 3h |
| 1-3 | +8h 30min | Fluorouracil (5-FU) | 10 $\mu\text{M}$ | 3h |
| *Dynamic dose FOLFIRI* |  |  |  |  |
| 1 | 0 | Irinotecan (SN38) | 50.7 nM | 2h |
| 1 | +2h | Irinotecan (SN38) | 50.7 nM | 30min |
| | | Fluorouracil (5-FU) | 94.5 $\mu\text{M}$ | |
| 1 | +2h 30min | Irinotecan (SN38) | 50.7 nM | 2h 30min |
| | | Fluorouracil (5-FU) | 3.4 $\mu\text{M}$ | |
| 1 | +5h | Irinotecan (SN38) | 21.1 nM | 18h 30min |
| | | Fluorouracil (5-FU) | 3.4 $\mu\text{M}$ | |
| 2 | 0 | Fluorouracil (5-FU) | 3.4 $\mu\text{M}$ | 26h |

### Sensitivity testing of colorectal cancer cells

#### The dynamic dose test better predicts the response to chemotherapy in CRC patients

At first, cells were exposed to a series of constant concentrations of the same drugs for 72 hours. The patients in this cohort (N=6) received several chemotherapy lines, including mFOLFOX-6, CapOx, FOLFIRI or Capecitabin mono schemes (more detailed information is provided in Table S1). Four patients did not respond to any treatment, while two responded to 5-FU–irinotecan combination therapy. The response values had a wide distribution with the 5-FU IC_50_ range = 30.25 to 984.37 *µ*M, oxaliplatin IC_50_ range = 29.88 to 294.66 *µ*M and SN-38 IC_50_ range = 33.55 to 103.99 nM. Surprisingly, the cells derived from the responder had 5-FU values on opposite sides of the spectrum (30.25 and 313.38 *µ*M). SN-38 IC_50_ values were on the lower end which is consistent with previous studies (10). However, it was not possible to determine a cutoff without misclassifying responders and non-responders. Therefore, the IC_50_ metrics did not prove to be effective for predicting the response to treatment in our study (Figure 6).

**Fig. 6.**
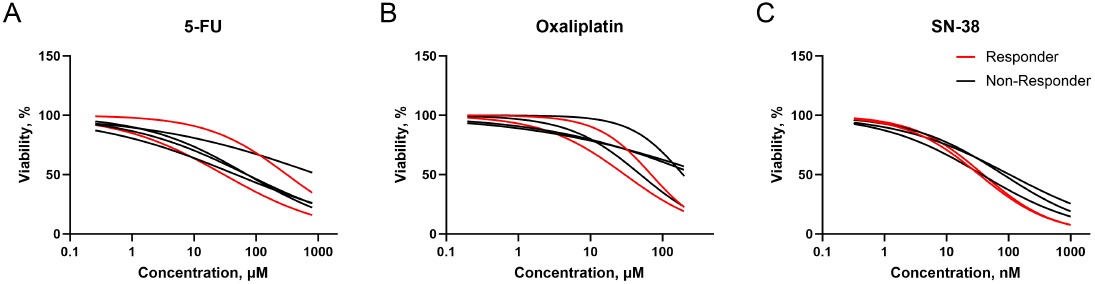
Fitted dose–response curves of the primary colorectal cancer cells exposed to the 5-FU (A), oxaliplatin (B) and SN-38 (C).

Similar findings were observed for the GI_50_ metrics (Figure 7). It was also not possible to identify quadrants in 2D of 5-FU - Oxaliplatin GI_50_ values (Figure 7D) and 5-FU - SN-38 GI_50_ values (Figure 7E) for the accurate stratification of cells based on the clinical response. In general, these results align with previous studies that suggest that the SN-38 IC_50_ or GI_50_ values rather than 5-FU are better biomarkers for predicting responses to the FOLFIRI scheme. Interestingly, we observed a stronger correlation between the IC_50_ and GI_50_ values for 5-FU and SN-38 than for oxaliplatin (Figure S1). Because the patients were treated with combination chemotherapy, we performed additional experiments with mixtures of the applied drugs. Cells were exposed to clinically reachable concentrations of 5-FU and oxaliplatin (3.4 *µ*M and 3.37 *µ*M) as well as 5-FU and SN-38 (3.4 *µ*M and 20.7 nM) combinations, or 5-FU alone (3.4 *µ*M) for 46 hours. The combination of 5-FU and oxaliplatin was used to assess response to mFOLFOX-6 or CapOx schemes, while 5-FU – SN-38 and 5-FU alone to evaluate responses to FOLFIRI or capecitabine monotherapy, respectively. The test results with clinically relevant concentrations made it possible to clearly identify non-responders to oxaliplatin-containing therapy schemes and capecitabine monotherapy (Figure 8A). These results were also more consistent with the clinical outcomes for the FOLFIRI scheme. Recently another study by Tan et al. Tan et al. (15) demonstrated a 83% accuracy in predicting responses in CRC patients supporting the importance of utilization of clinically relevant concentrations of chemotherapeutic drugs in *in vitro* tests. However, we were still unable to establish a cutoff value for the classification of responders and non-responders to irinotecan treatment using the IC_50_ and GI_50_ metrics. From a clinical utility perspective, the clear identification of resistance holds the same, if not higher, value as the prediction of sensitivity, because ineffective treatment cycles worsen a patient’s condition and limit further treatment options.

**Fig. 7.**
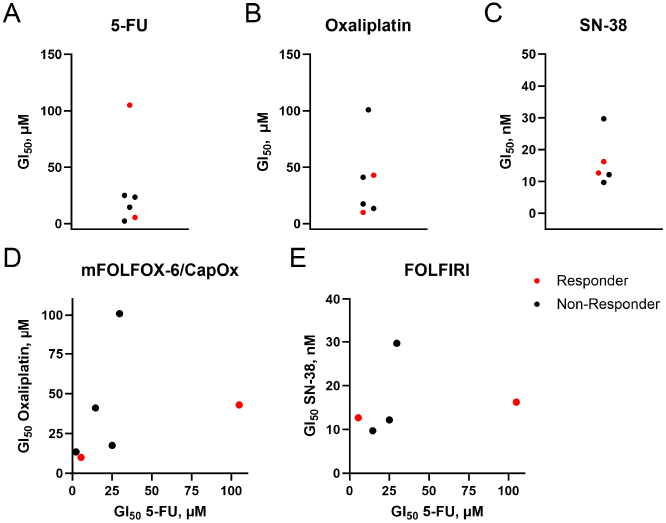
GI_50_ values of the primary colorectal cancer cells exposed to 5-FU (A), oxaliplatin (B), SN-38 (C) and 2D plots for the drugs associated with oxaliplatin (D) or irinotecan-containing schemes (E).

**Fig. 8.**
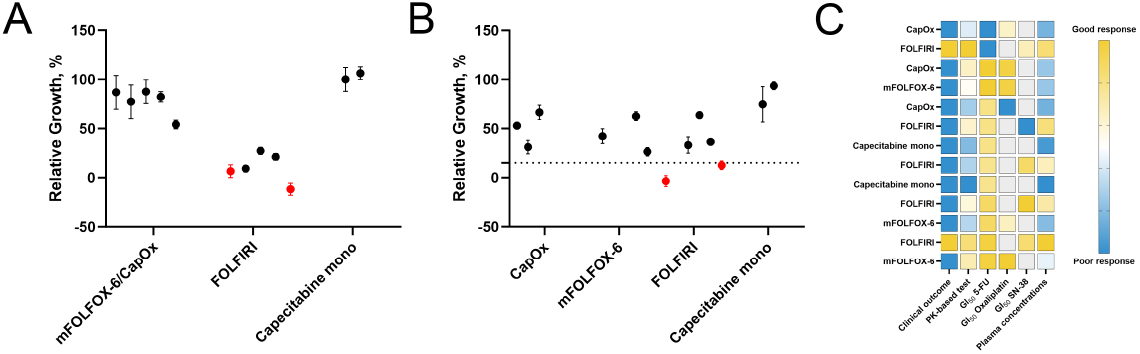
The *in vitro* response of primary colorectal cancer cells exposed to drug mixtures at clinically reachable concentrations (A), treated according to PK-based protocols (B), and a general comparison of responses using different methods (C).

Finally, to evaluate the performance of our *dynamic dose* test, we treated the cells according to the corresponding protocols for the FOLFOX-6, CapOx, and FOLFIRI schemes. As can be seen in (Figure 8B), shorter exposure times, recapitulation of the sequence and clinically relevant concentrations resulted in a more widely dispersed response distribution to the FOLFIRI treatment. Such an approach enabled stratification of the cells for all chemotherapy schemes tested using the cutoff of 15%. Therefore, the *dynamic dose* test demonstrated the best performance for classifying ‘responders’ and ‘non-responders’ in a small cohort of primary colorectal cancer cells among all the methods used (Figure 8C).

#### The dynamic dose test recapitulates the equivalence of the mFOLFOX-6 and CapOx

Another interesting result we observed during the evaluation of the response to mFOLFOX-6 and CapOx schemes on the primary colorectal cancer cells derived from patient 2. According to the generated *dynamic dose* mFOLFOX-6 and CapOx protocols (Table 2), cells were exposed to different concentrations of oxaliplatin (3.37 *µ*M for mFOLFOX-6 and 5.15 *µ*M for CapOx) and 5-FU (94.5 *µ*M followed by 3.4 *µ*M for mFOLFOX-6 and 10 *µ*M for CapOx) for different incubation times. This led to differences in AUC^PK^ for oxaliplatin (6.74 *µ*M·h for mFOLFOX-6 and 10.30 *µ*M·h for CapOx) and 5-FU (203.65 *µ*M·h for mFOLFOX-6 and 180.00 *µ*M·h for CapOx). Despite the differences in testing conditions, the response was similar for both schemes. Clinically, these two schemes have been demonstrated to exhibit no statistical differences in overall survival and overall response rate (74, 75). Therefore, we quantified the response of both primary colorectal cancer cells (Figure 9A) and CRC cell lines (Figure 9B) to evaluate whether the PK-based *dynamic dose* test could recapitulate these findings *in vitro*.

**Fig. 9.**
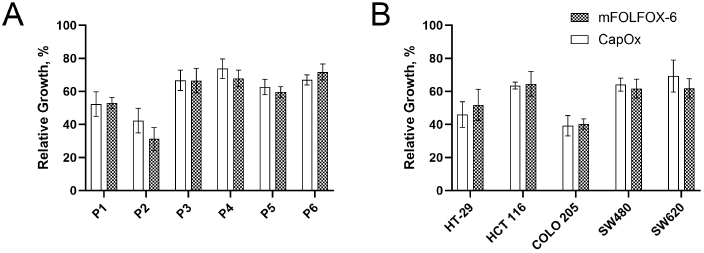
The *in vitro* response of primary colorectal cancer cells (A) and CRC cell lines (B) to mFOLFOX-6 and CapOx treatment.

As can be seen in (Figure 9), *dynamic dose* test results were consistent with the historical clinical data. *Dynamic dose tests* revealed a similar response of all colorectal cancer cells to mFOLFOX-6 and CapOx. This highlights the potential utility of this test for comparing the efficacy of treatment schemes during drug development.

## Conclusions

In 2021, the Oncology Center of Excellence has launched a Project Optimus initiative to reform the paradigm of dose optimization and dose selection in oncology drug development (76). The later published guidelines highlighted that the currently used clinical and *in vitro* approaches for dose finding (e.g. maximum tolerated dose) are not optimal and may not apply to the latest anticancer drugs, such as targeted therapies (77). Conventional drug testing methods under the static conditions that are used in early drug development may hamper the discovery of optimal drug doses. This initial non-optimal study design choice can lead to the need for further dose optimization studies in the late stages of clinical development resulting in a high failure rate and excessive costs. For example, sotorasib (78), capecitabine (79) and gemtuzumab (80) required post-approval dose adjustments to improve safety and tolerability. There are multiple dimensions for improving preclinical development, including more physiological disease modeling using organs-on-a-chip, development of more sophisticated cell models such as PDO and the utilization of computational multi-omics approaches. In our study, we have assessed the potential of pharmacokinetic-based *in vitro* testing to improve the drug sensitivity testing of colorectal cancer cells. The key results and their implications are outlined as follows:

- **Superior classification of ‘responders’ vs. ‘non-responders’:** Our PK-based *dynamic dose* test demonstrated the best performance for classifying ‘responders’ and ‘non-responders’ in a small cohort of primary colorectal cancer cells among all the methods used. The best result among classical approaches was achieved in a test with exposure to clinically reachable concentrations (only one misclassified case). IC_50_ and GI_50_ metrics were not univocal for different drugs and were particularly inaccurate for 5-FU.
- **Alignment with historical clinical data:** *Dynamic dose* test results were also consistent with the existing clinical data on similarities of mFOLFOX6 and CapOx schemes.
- **Application in drug development:** In addition to measuring individual patient responses, this approach also has the potential for ‘pharmacological calibration’ of a drug candidate before entering a phase I clinical trial (81). It includes optimising the dose and schedule to balance efficacy and toxicity, which current approaches are reported to underestimate (82).
- **Improvement of PK/PD modeling:** The dose-response function in PK/PD modeling is typically described by the Hill equation (83), which does not account for *in vitro* exposure time and may vary depending on the experimental setup. Integrating pharmacokinetics into *in vitro* tests enables direct measurement of dose-dependent drug effects, facilitating high-throughput data acquisition necessary for PK/PD modeling and optimizing schedules in the frequency domain (84).
- **Impact on personalized treatment:** One of the advantages of the *dynamic dose* test is its flexibility in adjusting the test procedure based on the treatment schedule and the individual PK characteristics of a drug, a patient, or a group of patients.

In sum, this research contributes to the ongoing effort to bridge the gap between *in vitro* assays and clinical outcomes. Although these assumptions have yet to be validated, the results presented in this study, along with findings from other groups, suggest that the PK-based dynamic dose testing approach may have considerable potential in fields of drug development and personalized treatment.

## Data availability

All additional data that supports the findings of this study is available on request from the corresponding author.

## Author contributions

A.A.P., B.R.B. - conceptualization, supervision; A.A.P., B.R.B., S.N. - data curation, formal analysis, investigation, methodology; A.A.P., B.R.B., S.R., K.-H.G., M.W., J.F. - original draft, review and editing.

## ACKNOWLEDGEMENTS

We thank Prof. Udo Schumacher, Dr. Jens Hoffmann and Dr. Martin Schumacher for their valuable recommendations regarding methodology and manuscript structure.

## Supplementary material

**Table S1.** Treatment history of the patients.

| Patient ID | Initial Diagnosis | Treatment | Clinical outcome |
| --- | --- | --- | --- |
| 1 | pT4N1M0 | XELOX | Progressive disease |
|  |  | FOLFIRI | Partial response |
| 2 | pT4aN2M0 | XELOX | Progressive disease |
|  |  | FOLFOX | Progressive disease |
| 3 | pT2N1aM0 | XELOX | Progressive disease |
|  |  | FOLFIRI | Progressive disease |
|  |  | Capecitabine mono | Progressive disease |
| 4 | cT3N2aM1 | FOLFIRI | Progressive disease |
|  |  | Capecitabine mono | Progressive disease |
| 5 | T4aN1M1 | FOLFIRI | Progressive disease |
|  |  | FOLFOX | Progressive disease |
| 6 | pT3N1aM1 | FOLFOX | Progressive disease |
|  |  | FOLFIRI | Partial response |

**Fig. S1.**
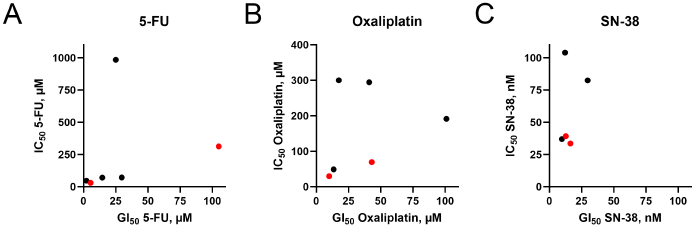
Correlation between IC_50_ and GI_50_ for the drugs tested.

## Bibliography

1. J. L. Sebaugh. Guidelines for accurate EC50/IC50 estimation. Pharmaceutical Statistics, 10(2):128–134, March 2011. ISSN 1539-1604, 1539-1612. doi: 10.1002/pst.426.

2. Jordi Barretina, Giordano Caponigro, Nicolas Stransky, Kavitha Venkatesan, Adam A. Margolin, Sungjoon Kim, Christopher J. Wilson, Joseph Lehár, Gregory V. Kryukov, Dmitriy Sonkin, Anupama Reddy, Manway Liu, Lauren Murray, Michael F. Berger, John E. Monahan, Paula Morais, Jodi Meltzer, Adam Korejwa, Judit Jané-Valbuena, Felipa A. Mapa, Joseph Thibault, Eva Bric-Furlong, Pichai Raman, Aaron Shipway, Ingo H. Engels, Jill Cheng, Guoying K. Yu, Jianjun Yu, Peter Aspesi, Melanie De Silva, Kalpana Jagtap, Michael D. Jones, Li Wang, Charles Hatton, Emanuele Palescandolo, Supriya Gupta, Scott Mahan, Carrie Sougnez, Robert C. Onofrio, Ted Liefeld, Laura MacConaill, Wendy Winckler, Michael Reich, Nanxin Li, Jill P. Mesirov, Stacey B. Gabriel, Gad Getz, Kristin Ardlie, Vivien Chan, Vic E. Myer, Barbara L. Weber, Jeff Porter, Markus Warmuth, Peter Finan, Jennifer L. Harris, Matthew Meyerson, Todd R. Golub, Michael P. Morrissey, William R. Sellers, Robert Schlegel, and Levi A. Garraway. The Cancer Cell Line Encyclopedia enables predictive modelling of anticancer drug sensitivity. Nature, 483(7391):603–607, March 2012. ISSN 0028-0836, 1476-4687. doi: 10.1038/nature11003.

3. Mahmoud Ghandi, Franklin W. Huang, Judit Jané-Valbuena, Gregory V. Kryukov, Christopher C. Lo, E. Robert McDonald, Jordi Barretina, Ellen T. Gelfand, Craig M. Bielski, Haoxin Li, Kevin Hu, Alexander Y. Andreev-Drakhlin, Jaegil Kim, Julian M. Hess, Brian J. Haas, François Aguet, Barbara A. Weir, Michael V. Rothberg, Brenton R. Paolella, Michael S. Lawrence, Rehan Akbani, Yiling Lu, Hong L. Tiv, Prafulla C. Gokhale, Antoine De Weck, Ali Amin Mansour, Coyin Oh, Juliann Shih, Kevin Hadi, Yanay Rosen, Jonathan Bistline, Kavitha Venkatesan, Anupama Reddy, Dmitriy Sonkin, Manway Liu, Joseph Lehar, Joshua M. Korn, Dale A. Porter, Michael D. Jones, Javad Golji, Giordano Caponigro, Jordan E. Taylor, Caitlin M. Dunning, Amanda L. Creech, Allison C. Warren, James M. McFarland, Mahdi Zamanighomi, Audrey Kauffmann, Nicolas Stransky, Marcin Imielinski, Yosef E. Maruvka, Andrew D. Cherniack, Aviad Tsherniak, Francisca Vazquez, Jacob D. Jaffe, Andrew A. Lane, David M. Weinstock, Cory M. Johannessen, Michael P. Morrissey, Frank Stegmeier, Robert Schlegel, William C. Hahn, Gad Getz, Gordon B. Mills, Jesse S. Boehm, Todd R. Golub, Levi A. Garraway, and William R. Sellers. Next-generation characterization of the Cancer Cell Line Encyclopedia. Nature, 569(7757):503–508, May 2019. ISSN 0028-0836, 1476-4687. doi: 10.1038/s41586-019-1186-3.

4. James Inglese, Douglas S. Auld, Ajit Jadhav, Ronald L. Johnson, Anton Simeonov, Adam Yasgar, Wei Zheng, and Christopher P. Austin. Quantitative high-throughput screening: A titration-based approach that efficiently identifies biological activities in large chemical libraries. Proceedings of the National Academy of Sciences, 103(31):11473–11478, August 2006. ISSN 0027-8424, 1091-6490. doi: 10.1073/pnas.0604348103.

5. Natalie De Souza. Organoids. Nature Methods, 15(1):23–23, January 2018. ISSN 1548-7091, 1548-7105. doi: 10.1038/nmeth.4576.

6. Yuting Tang, Ting Wang, Yaowen Hu, Hongli Ji, Botao Yan, Xiarong Hu, Yunli Zeng, Yifan Hao, Weisong Xue, Zexin Chen, Jianqiang Lan, Yanan Wang, Haijun Deng, Chuxia Deng, Xiufeng Wu, and Jun Yan. Cutoff value of IC50 for drug sensitivity in patient-derived tumor organoids in colorectal cancer. iScience, 26(7):107116, July 2023. ISSN 25890042. doi: 10.1016/j.isci.2023.107116.

7. G. Emerens Wensink, Sjoerd G. Elias, Jasper Mullenders, Miriam Koopman, Sylvia F. Boj, Onno W. Kranenburg, and Jeanine M. L. Roodhart. Patient-derived organoids as a predictive biomarker for treatment response in cancer patients. npj Precision Oncology, 5(1):30, April 2021. ISSN 2397-768X. doi: 10.1038/s41698-021-00168-1.

8. Karuna Ganesh, Chao Wu, Kevin P. O’Rourke, Bryan C. Szeglin, Youyun Zheng, Charles-Etienne Gabriel Sauvé, Mohammad Adileh, Isaac Wasserman, Michael R. Marco, Amanda S. Kim, Maha Shady, Francisco Sanchez-Vega, Wouter R. Karthaus, Helen H. Won, Seo-Hyun Choi, Raphael Pelossof, Afsar Barlas, Peter Ntiamoah, Emmanouil Pappou, Arthur Elghouayel, James S. Strong, Chin-Tung Chen, Jennifer W. Harris, Martin R. Weiser, Garrett M. Nash, Jose G. Guillem, Iris H. Wei, Richard N. Kolesnick, Harini Veeraraghavan, Eduardo J. Ortiz, Iva Petkovska, Andrea Cercek, Katia O. Manova-Todorova, Leonard B. Saltz, Jessica A. Lavery, Ronald P. DeMatteo, Joan Massagué, Philip B. Paty, Rona Yaeger, Xi Chen, Sujata Patil, Hans Clevers, Michael F. Berger, Scott W. Lowe, Jinru Shia, Paul B. Romesser, Lukas E. Dow, Julio Garcia-Aguilar, Charles L. Sawyers, and J. Joshua Smith. A rectal cancer organoid platform to study individual responses to chemoradiation. Nature Medicine, 25(10):1607–1614, October 2019. ISSN 1078-8956, 1546-170X. doi: 10.1038/s41591-019-0584-2.

9. Vignesh Narasimhan, Josephine A. Wright, Michael Churchill, Tongtong Wang, Rachele Rosati, Tamsin R.M. Lannagan, Laura Vrbanac, Anne B. Richardson, Hiroki Kobayashi, Timothy Price, Gayle X.Y. Tye, Julie Marker, Peter J. Hewett, Michael P. Flood, Shalini Pereira, G. Adam Whitney, Michael Michael, Jeanne Tie, Siddhartha Mukherjee, Carla Grandori, Alexander G. Heriot, Daniel L. Worthley, Robert G. Ramsay, and Susan L. Woods. Medium-throughput Drug Screening of Patient-derived Organoids from Colorectal Peritoneal Metastases to Direct Personalized Therapy. Clinical Cancer Research, 26(14):3662–3670, July 2020. ISSN 1078-0432, 1557-3265. doi: 10.1158/1078-0432.CCR-20-0073.

10. Salo N. Ooft, Fleur Weeber, Krijn K. Dijkstra, Chelsea M. McLean, Sovann Kaing, Erik van Werkhoven, Luuk Schipper, Louisa Hoes, Daniel J. Vis, Joris van de Haar, Warner Prevoo, Petur Snaebjornsson, Daphne van der Velden, Michelle Klein, Myriam Chalabi, Henk Boot, Monique van Leerdam, Haiko J. Bloemendal, Laurens V. Beerepoot, Lodewyk Wessels, Edwin Cuppen, Hans Clevers, and Emile E. Voest. Patient-derived organoids can predict response to chemotherapy in metastatic colorectal cancer patients. Science Translational Medicine, 11(513):eaay2574, October 2019. doi: 10.1126/scitranslmed.aay2574. Publisher: American Association for the Advancement of Science.

11. Hervé Tiriac Pascal Belleau, Dannielle D. Engle, Dennis Plenker, Astrid Deschênes, Tim D. D. Somerville, Fieke E. M. Froeling, Richard A. Burkhart, Robert E. Denroche, Gun-Ho Jang, Koji Miyabayashi, C. Megan Young, Hardik Patel, Michelle Ma, Joseph F. LaComb, Randze Lerie D. Palmaira, Ammar A. Javed, Jasmine C. Huynh, Molly Johnson, Kanika Arora, Nicolas Robine, Minita Shah, Rashesh Sanghvi, Austin B. Goetz, Cinthya Y. Lowder, Laura Martello, Else Driehuis, Nicolas LeComte, Gokce Askan, Christine A. Iacobuzio-Donahue, Hans Clevers, Laura D. Wood, Ralph H. Hruban, Elizabeth Thompson, Andrew J. Aguirre, Brian M. Wolpin, Aaron Sasson, Joseph Kim, Maoxin Wu, Juan Carlos Bucobo, Peter Allen, Divyesh V. Sejpal, William Nealon, James D. Sullivan, Jordan M. Winter, Phyllis A. Gimotty, Jean L. Grem, Dominick J. DiMaio, Jonathan M. Buscaglia, Paul M. Grandgenett, Jonathan R. Brody, Michael A. Hollingsworth, Grainne M. O’Kane, Faiyaz Notta, Edward Kim, James M. Crawford, Craig Devoe, Allyson Ocean, Christopher L. Wolfgang, Kenneth H. Yu, Ellen Li, Christopher R. Vakoc, Benjamin Hubert, Sandra E. Fischer, Julie M. Wilson, Richard Moffitt, Jennifer Knox, Alexander Krasnitz, Steven Gallinger, and David A. Tuveson. Organoid Profiling Identifies Common Responders to Chemotherapy in Pancreatic Cancer. Cancer Discovery, 8(9):1112–1129, September 2018. ISSN 2159-8274, 2159-8290. doi: 10.1158/2159-8290.CD-18-0349.

12. Shuguang Huang and Lei Pang. Comparing Statistical Methods for Quantifying Drug Sensitivity Based on In Vitro Dose–Response Assays. ASSAY and Drug Development Technologies, 10(1):88–96, February 2012. ISSN 1540-658X, 1557-8127. doi: 10.1089/adt.2011.0388.

13. Marc Hafner, Mario Niepel, Mirra Chung, and Peter K Sorger. Growth rate inhibition metrics correct for confounders in measuring sensitivity to cancer drugs. Nature Methods, 13(6): 521–527, June 2016. ISSN 1548-7091, 1548-7105. doi: 10.1038/nmeth.3853.

14. Marc Hafner, Mario Niepel, and Peter K Sorger. Alternative drug sensitivity metrics improve preclinical cancer pharmacogenomics. Nature Biotechnology, 35(6):500–502, June 2017. ISSN 1087-0156, 1546-1696. doi: 10.1038/nbt.3882.

15. Tao Tan, Dmitri Mouradov, Margaret Lee, Grace Gard, Yumiko Hirokawa, Shan Li, Cong Lin, Fuqiang Li, Huijuan Luo, Kui Wu, Michelle Palmieri, Evelyn Leong, Jordan Clarke, Anuratha Sakthianandeswaren, Helen Brasier, Jeanne Tie, Niall C. Tebbutt, Azim Jalali, Rachel Wong, Antony W. Burgess, Peter Gibbs, and Oliver M. Sieber. Unified framework for patient-derived, tumor-organoid-based predictive testing of standard-of-care therapies in metastatic colorectal cancer. Cell Reports Medicine, 4(12):101335, December 2023. ISSN 26663791. doi: 10.1016/j.xcrm.2023.101335.

16. Tuomo Kalliokoski, Christian Kramer, Anna Vulpetti, and Peter Gedeck. Comparability of Mixed IC50 Data – A Statistical Analysis. PLoS ONE, 8(4):e61007, April 2013. ISSN 1932-6203. doi: 10.1371/journal.pone.0061007.

17. Peter Larsson, Hanna Engqvist, Jana Biermann, Elisabeth Werner Rönnerman, Eva Forssell-Aronsson, Anikó Kovács, Per Karlsson, Khalil Helou, and Toshima Z. Parris. Optimization of cell viability assays to improve replicability and reproducibility of cancer drug sensitivity screens. Scientific Reports, 10(1):5798, April 2020. ISSN 2045-2322. doi: 10.1038/s41598-020-62848-5.

18. Elin Jerremalm, Mikael Hedeland, Inger Wallin, Ulf Bondesson, and Hans Ehrsson. Oxaliplatin Degradation in the Presence of Chloride: Identification and Cytotoxicity of the Monochloro Monooxalato Complex. Pharmaceutical Research, 21(5):891–894, May 2004. ISSN 0724-8741. doi: 10.1023/B:PHAM.0000026444.67883.83.

19. Dane R. Liston and Myrtle Davis. Clinically Relevant Concentrations of Anticancer Drugs: A Guide for Nonclinical Studies. Clinical Cancer Research, 23(14):3489–3498, July 2017. ISSN 1078-0432, 1557-3265. doi: 10.1158/1078-0432.CCR-16-3083.

20. David M. Evans, Jianwen Fang, Thomas Silvers, Rene Delosh, Julie Laudeman, Chad Ogle, Russell Reinhart, Michael Selby, Lori Bowles, John Connelly, Erik Harris, Julia Krushkal, Larry Rubinstein, James H. Doroshow, and Beverly A. Teicher. Exposure time versus cytotoxicity for anticancer agents. Cancer Chemotherapy and Pharmacology, 84(2):359–371, August 2019. ISSN 0344-5704, 1432-0843. doi: 10.1007/s00280-019-03863-w.

21. Joe T. Sharick, Christine M. Walsh, Carley M. Sprackling, Cheri A. Pasch, Dan L. Pham, Karla Esbona, Alka Choudhary, Rebeca Garcia-Valera, Mark E. Burkard, Stephanie M. McGregor, Kristina A. Matkowskyj, Alexander A. Parikh, Ingrid M. Meszoely, Mark C. Kelley, Susan Tsai, Dustin A. Deming, and Melissa C. Skala. Metabolic Heterogeneity in Patient Tumor-Derived Organoids by Primary Site and Drug Treatment. Frontiers in Oncology, 10: 553, May 2020. ISSN 2234-943X. doi: 10.3389/fonc.2020.00553.

22. Georgios Vlachogiannis, Somaieh Hedayat, Alexandra Vatsiou, Yann Jamin, Javier Fernández-Mateos, Khurum Khan, Andrea Lampis, Katherine Eason, Ian Huntingford, Rosemary Burke, Mihaela Rata, Dow-Mu Koh, Nina Tunariu, David Collins, Sanna Hulkki-Wilson, Chanthirika Ragulan, Inmaculada Spiteri, Sing Yu Moorcraft, Ian Chau, Sheela Rao, David Watkins, Nicos Fotiadis, Maria Bali, Mahnaz Darvish-Damavandi, Hazel Lote, Zakaria Eltahir, Elizabeth C. Smyth, Ruwaida Begum, Paul A. Clarke, Jens C. Hahne, Mitchell Dowsett, Johann De Bono, Paul Workman, Anguraj Sadanandam, Matteo Fassan, Owen J. Sansom, Suzanne Eccles, Naureen Starling, Chiara Braconi, Andrea Sottoriva, Simon P. Robinson, David Cunningham, and Nicola Valeri. Patient-derived organoids model treatment response of metastatic gastrointestinal cancers. Science, 359(6378):920–926, February 2018. ISSN 0036-8075, 1095-9203. doi: 10.1126/science.aao2774.

23. Sangmin Choe and Donghwan Lee. Parameter estimation for sigmoid E _max_ models in exposure-response relationship. Translational and Clinical Pharmacology, 25(2):74, 2017. ISSN 2289-0882, 2383-5427. doi: 10.12793/tcp.2017.25.2.74.

24. N. Srinivas. Dual Incorporation of the in vitro Data (IC50) and in vivo (Cmax) Data for the Prediction of Area Under the Curve (AUC) for Statins using Regression Models Developed for Either Pravastatin or Simvastatin. Drug Research, 66(08):402–406, May 2016. ISSN 2194-9379, 2194-9387. doi: 10.1055/s-0042-106289.

25. Monika Engelhardt, Roland Mertelsmann, and Justus Duyster, editors. Das Blaue Buch: Chemotherapie-Manual Hämatologie und Onkologie. Springer Berlin Heidelberg, Berlin, Heidelberg, 2020. ISBN 978-3-662-60379-6 978-3-662-60380-2. doi: 10.1007/978-3-662-60380-2.

26. Ting-Chao Chou. Drug Combination Studies and Their Synergy Quantification Using the Chou-Talalay Method. Cancer Research, 70(2):440–446, January 2010. ISSN 0008-5472, 1538-7445. doi: 10.1158/0008-5472.CAN-09-1947.

27. Ting-Chao Chou. Preclinical versus clinical drug combination studies. Leukemia & Lymphoma, 49(11):2059–2080, January 2008. ISSN 1042-8194, 1029-2403. doi: 10.1080/10428190802353591.

28. J-L Fischel, P Rostagno, P Formento, A Dubreuil, M-C Etienne, and G Milano. Ternary combination of irinotecan, fluorouracil-folinic acid and oxaliplatin: results on human colon cancer cell lines. British Journal of Cancer, 84(4):579–585, February 2001. ISSN 15321827. doi: 10.1054/bjoc.2000.1600.

29. Johan Gabrielsson, Lambertus A. Peletier, and Stephan Hjorth. Lost in translation: What’s in an EC? Innovative PK/PD reasoning in the drug development context. European Journal of Pharmacology, 835:154–161, September 2018. ISSN 00142999. doi: 10.1016/j.ejphar.2018.07.037.

30. Rasmus Jansson-Löfmark, Stephan Hjorth, and Johan Gabrielsson. Does In Vitro Potency Predict Clinically Efficacious Concentrations? Clinical Pharmacology & Therapeutics, 108 (2):298–305, August 2020. ISSN 0009-9236, 1532-6535. doi: 10.1002/cpt.1846.

31. Giuseppe Pizzorno and Robert E. Handschumacher. Effect of clinically modeled regimens on the growth response and development of resistance in human colon carcinoma cell lines. Biochemical Pharmacology, 49(4):559–565, February 1995. ISSN 00062952. doi: 10.1016/0006-2952(94)00445-R.

32. Patrick Erickson, Gunjan Jetley, Param Amin, Aamena Mejevdiwala, Ashna Patel, Kelli Cheng, and Biju Parekkadan. A cell culture system to model pharmacokinetics using adjustable-volume perfused mixing chambers. Toxicology in Vitro, 91:105623, September 2023. ISSN 08872333. doi: 10.1016/j.tiv.2023.105623.

33. Christian Lohasz, Jacqueline Loretan, Dario Sterker, Ekkehard Görlach, Kasper Renggli, Paul Argast, Olivier Frey, Marion Wiesmann, Markus Wartmann, Martin Rausch, and Andreas Hierlemann. A Microphysiological Cell-Culturing System for Pharmacokinetic Drug Exposure and High-Resolution Imaging of Arrays of 3D Microtissues. Frontiers in Pharmacology, 12:785851, December 2021. ISSN 1663-9812. doi: 10.3389/fphar.2021.785851.

34. Christian Lohasz, Olivier Frey, Flavio Bonanini, Kasper Renggli, and Andreas Hierlemann. Tubing-Free Microfluidic Microtissue Culture System Featuring Gradual, in vivo-Like Substance Exposure Profiles. Frontiers in Bioengineering and Biotechnology, 7:72, April 2019. ISSN 2296-4185. doi: 10.3389/fbioe.2019.00072.

35. Job Komen, Eiko Y. Westerbeek, Ruben W. Kolkman, Julia Roesthuis, Caroline Lievens, Albert Van Den Berg, and Andries D. Van Der Meer. Controlled pharmacokinetic anti-cancer drug concentration profiles lead to growth inhibition of colorectal cancer cells in a microfluidic device. Lab on a Chip, 20(17):3167–3178, 2020. ISSN 1473-0197, 1473-0189. doi: 10.1039/D0LC00419G.

36. Tudor Petreus, Elaine Cadogan, Gareth Hughes, Aaron Smith, Venkatesh Pilla Reddy, Alan Lau, Mark James O’Connor, Susan Critchlow, Marianne Ashford, and Lenka Oplustil O’Connor. Tumour-on-chip microfluidic platform for assessment of drug pharmacokinetics and treatment response. Communications Biology, 4(1):1001, August 2021. ISSN 2399-3642. doi: 10.1038/s42003-021-02526-y.

37. Dharaminder Singh, Sudhir P. Deosarkar, Elaine Cadogan, Vikki Flemington, Alysha Bray, Jingwen Zhang, Ronald S. Reiserer, David K. Schaffer, Gregory B. Gerken, Clayton M. Britt, Erik M. Werner, Francis D. Gibbons, Tomasz Kostrzewski, Christopher E. Chambers, Emma J. Davies, Antonio Ramos Montoya, Jacqueline H. L. Fok, David Hughes, Kristin Fabre, Matthew P. Wagoner, John P. Wikswo, and Clay W. Scott. A microfluidic system that replicates pharmacokinetic (PK) profiles in vitro improves prediction of in vivo efficacy in preclinical models. PLOS Biology, 20(5):e3001624, May 2022. ISSN 1545-7885. doi: 10.1371/journal.pbio.3001624.

38. Myriam Chalabi, Lorenzo F. Fanchi, Krijn K. Dijkstra, José G. Van Den Berg, Arend G. Aalbers, Karolina Sikorska, Marta Lopez-Yurda, Cecile Grootscholten, Geerard L. Beets, Petur Snaebjornsson, Monique Maas, Marjolijn Mertz, Vivien Veninga, Gergana Bounova, Annegien Broeks, Regina G. Beets-Tan, Thomas R. De Wijkerslooth, Anja U. Van Lent, Hendrik A. Marsman, Elvira Nuijten, Niels F. Kok, Maria Kuiper, Wieke H. Verbeek, Marleen Kok, Monique E. Van Leerdam, Ton N. Schumacher, Emile E. Voest, and John B. Haanen. Neoadjuvant immunotherapy leads to pathological responses in MMR-proficient and MMR-deficient early-stage colon cancers. Nature Medicine, 26(4):566–576, April 2020. ISSN 1078-8956, 1546-170X. doi: 10.1038/s41591-020-0805-8.

39. Ye Yao, Xiaoya Xu, Lifeng Yang, Ji Zhu, Juefeng Wan, Lijun Shen, Fan Xia, Guoxiang Fu, Yun Deng, Mengxue Pan, Qiang Guo, Xiaoxue Gao, Yuanchuang Li, Xinxin Rao, Yi Zhou, Liping Liang, Yaqi Wang, Jing Zhang, Hui Zhang, Guichao Li, Lixing Zhang, Junjie Peng, Sanjun Cai, Chen Hu, Jianjun Gao, Hans Clevers, Zhen Zhang, and Guoqiang Hua. Patient-Derived Organoids Predict Chemoradiation Responses of Locally Advanced Rectal Cancer. Cell Stem Cell, 26(1):17–26.e6, January 2020. ISSN 19345909. doi: 10.1016/j.stem.2019.10.010.

40. Nina G. Steele, Jayati Chakrabarti, Jiang Wang, Jacek Biesiada, Loryn Holokai, Julie Chang, Lauren M. Nowacki, Jennifer Hawkins, Maxime Mahe, Nambirajan Sundaram, Noah Shroyer, Mario Medvedovic, Michael Helmrath, Syed Ahmad, and Yana Zavros. An Organoid-Based Preclinical Model of Human Gastric Cancer. Cellular and Molecular Gastroenterology and Hepatology, 7(1):161–184, 2019. ISSN 2352345X. doi: 10.1016/j.jcmgh.2018.09.008.

41. Norman Sachs, Joep De Ligt, Oded Kopper, Ewa Gogola, Gergana Bounova, Fleur Weeber, Anjali Vanita Balgobind, Karin Wind, Ana Gracanin, Harry Begthel, Jeroen Korving, Ruben Van Boxtel, Alexandra Alves Duarte, Daphne Lelieveld, Arne Van Hoeck, Robert Frans Ernst, Francis Blokzijl, Isaac Johannes Nijman, Marlous Hoogstraat, Marieke Van De Ven, David Anthony Egan, Vittoria Zinzalla, Jurgen Moll, Sylvia Fernandez Boj, Emile Eugene Voest, Lodewyk Wessels, Paul Joannes Van Diest, Sven Rottenberg, Robert Gerhardus Jacob Vries, Edwin Cuppen, and Hans Clevers. A Living Biobank of Breast Cancer Organoids Captures Disease Heterogeneity. Cell, 172(1-2):373–386.e10, January 2018. ISSN 00928674. doi: 10.1016/j.cell.2017.11.010.

42. Chris Jenske De Witte, Jose Espejo Valle-Inclan, Nizar Hami, Kadi Lõhmussaar, Oded Kopper, Celien Philomena Henrieke Vreuls, Geertruida Nellie Jonges, Paul Van Diest, Luan Nguyen, Hans Clevers, Wigard Pieter Kloosterman, Edwin Cuppen, Hugo Johannes Gerhardus Snippert, Ronald Peter Zweemer, Petronella Oda Witteveen, and Ellen Stelloo. Patient-Derived Ovarian Cancer Organoids Mimic Clinical Response and Exhibit Heterogeneous Inter- and Intrapatient Drug Responses. Cell Reports, 31(11):107762, June 2020. ISSN 22111247. doi: 10.1016/j.celrep.2020.107762.

43. Nhan Phan, Jenny J. Hong, Bobby Tofig, Matthew Mapua, David Elashoff, Neda A. Moatamed, Jin Huang, Sanaz Memarzadeh, Robert Damoiseaux, and Alice Soragni. A simple high-throughput approach identifies actionable drug sensitivities in patient-derived tumor organoids. Communications Biology, 2(1):78, February 2019. ISSN 2399-3642. doi: 10.1038/s42003-019-0305-x.

44. Konstantinos I. Votanopoulos, Steven Forsythe, Hemamylammal Sivakumar, Andrea Mazzocchi, Julio Aleman, Lance Miller, Edward Levine, Pierre Triozzi, and Aleksander Skardal. Model of Patient-Specific Immune-Enhanced Organoids for Immunotherapy Screening: Feasibility Study. Annals of Surgical Oncology, 27(6):1956–1967, June 2020. ISSN 1068-9265, 1534-4681. doi: 10.1245/s10434-019-08143-8.

45. Xiaodun Li, Hayley E. Francies, Maria Secrier, Juliane Perner, Ahmad Miremadi, Núria Galeano-Dalmau, William J. Barendt, Laura Letchford, Genevieve M. Leyden, Emma K. Goffin, Andrew Barthorpe, Howard Lightfoot, Elisabeth Chen, James Gilbert, Ayesha Noorani, Ginny Devonshire, Lawrence Bower, Amber Grantham, Shona MacRae, Nicola Grehan, David C. Wedge, Rebecca C. Fitzgerald, and Mathew J. Garnett. Organoid cultures recapitulate esophageal adenocarcinoma heterogeneity providing a model for clonality studies and precision therapeutics. Nature Communications, 9(1):2983, July 2018. ISSN 2041-1723. doi: 10.1038/s41467-018-05190-9.

46. Andrea R. Mazzocchi, Shiny A. P. Rajan, Konstantinos I. Votanopoulos, Adam R. Hall, and Aleksander Skardal. In vitro patient-derived 3D mesothelioma tumor organoids facilitate patient-centric therapeutic screening. Scientific Reports, 8(1):2886, February 2018. ISSN 2045-2322. doi: 10.1038/s41598-018-21200-8.

47. Andrey Poloznikov, Sergey Nikulin, Larisa Bolotina, Andrei Kachmazov, Maria Raigorodskaya, Anna Kudryavtseva, Ildar Bakhtogarimov, Sergey Rodin, Irina Gaisina, Maxim Topchiy, Andrey Asachenko, Victor Novosad, Alexander Tonevitsky, and Boris Alekseev. 9-ING-41, a Small Molecule Inhibitor of GSK-3β, Potentiates the Effects of Chemotherapy on Colorectal Cancer Cells. Frontiers in Pharmacology, 12:777114, December 2021. ISSN 1663-9812. doi: 10.3389/fphar.2021.777114.

48. Alexandra Razumovskaya, Mariia Silkina, Andrey Poloznikov, Timur Kulagin, Maria Raigorodskaya, Nina Gorban, Anna Kudryavtseva, Maria Fedorova, Boris Alekseev, Alexander Tonevitsky, and Sergey Nikulin. Predicting patient outcomes with gene-expression biomarkers from colorectal cancer organoids and cell lines. Frontiers in Molecular Biosciences, Volume 12 - 2025, 2025. ISSN 2296-889X. doi: 10.3389/fmolb.2025.1531175.

49. Per-Anders Larsson, Göran Carlsson, Bengt Gustavsson, Wilhelm Graf, and Bengt Glimelius. Different Intravenous Administration Techniques for 5-Fluorouracil Pharmacokinetics and Pharmacodynamic Effects. Acta Oncologica, 35(2):207–212, January 1996. ISSN 0284-186X, 1651-226X. doi: 10.3109/02841869609098503.

50. M. A. Graham, G. F. Lockwood, D. Greenslade, S. Brienza, M. Bayssas, and E. Gamelin. Clinical pharmacokinetics of oxaliplatin: a critical review. Clinical Cancer Research: An Official Journal of the American Association for Cancer Research, 6(4):1205–1218, April 2000. ISSN 1078-0432.

51. Taroh Satoh, Hirofumi Yasui, Kei Muro, Yoshito Komatsu, Shinichi Sameshima, Kensei Yamaguchi, and Kenichi Sugihara. Pharmacokinetic assessment of irinotecan, SN-38, and SN-38-glucuronide: a substudy of the FIRIS study. Anticancer Research, 33(9):3845–3853, September 2013. ISSN 1791-7530.

52. Matthew D. Hall, Katherine A. Telma, Ki-Eun Chang, Tobie D. Lee, James P. Madigan, John R. Lloyd, Ian S. Goldlust, James D. Hoeschele, and Michael M. Gottesman. Say No to DMSO: Dimethylsulfoxide Inactivates Cisplatin, Carboplatin, and Other Platinum Complexes. Cancer Research, 74(14):3913–3922, July 2014. ISSN 0008-5472, 1538-7445. doi: 10.1158/0008-5472.CAN-14-0247.

53. R Core Team. R: A Language and Environment for Statistical Computing. R Foundation for Statistical Computing, Vienna, Austria, 2024.

54. Posit team. RStudio: Integrated Development Environment for R. Posit Software, PBC, Boston, MA, 2024.

55. Hadley Wickham, Romain François, Lionel Henry, Kirill Müller, and Davis Vaughan. dplyr: A Grammar of Data Manipulation, 2023.

56. Hadley Wickham, Davis Vaughan, and Maximilian Girlich. tidyr: Tidy Messy Data, 2024.

57. Hadley Wickham. ggplot2: Elegant Graphics for Data Analysis. Springer-Verlag New York, 2016. ISBN 978-3-319-24277-4.

58. Nan Xiao. ggsci: Scientific Journal and Sci-Fi Themed Color Palettes for ‘ggplot2’, 2023. R package version 3.2.0, https://github.com/nanxstats/ggsci.

59. Teun van den Brand. ggh4x: Hacks for ‘ggplot2’, 2023. R package version 1.3.1.

60. Thomas Lin Pedersen. patchwork: The Composer of Plots, 2024. R package version 1.3.0.9000, https://github.com/thomasp85/patchwork.

61. Zhexue Huang. Clustering Large Data Sets with Mixed Numeric and Categorical Values. In Proceedings of the First Pacific Asia Knowledge Discovery and Data Mining Conference, pages 21–34, Singapore, June 1997.

62. Zhexue Huang. Extensions to the k-Means Algorithm for Clustering Large Data Sets with Categorical Values. Data Mining and Knowledge Discovery, 2(3):283–304, 1998. ISSN 13845810. doi: 10.1023/A:1009769707641.

63. Francis Lévi, Gerard Metzger, Claire Massari, and Gerard Milano. Oxaliplatin: Pharmacokinetics and Chronopharmacological Aspects. Clinical Pharmacokinetics, 38(1):1–21, January 2000. ISSN 0312-5963. doi: 10.2165/00003088-200038010-00001.

64. Elin Jerremalm, Inger Wallin, and Hans Ehrsson. New insights into the biotransformation and pharmacokinetics of oxaliplatin. Journal of Pharmaceutical Sciences, 98(11):3879–3885, November 2009. ISSN 00223549. doi: 10.1002/jps.21732.

65. H. Ehrsson, I. Wallin, and J. Yachnin. Pharmacokinetics of Oxaliplatin in Humans. Medical Oncology, 19(4):261–266, 2002. ISSN 1357-0560. doi: 10.1385/MO:19:4:261.

66. Hans Ehrsson and Inger Wallin. Liquid chromatographic determination of oxaliplatin in blood using post-column derivatization in a microwave field followed by photometric detection. Journal of Chromatography B, 795(2):291–294, October 2003. ISSN 15700232. doi: 10.1016/S1570-0232(03)00590-7.

67. G. Bocci, R. Danesi, A. D. Di Paolo, F. Innocenti, G. Allegrini, A. Falcone, A. Melosi, M. Battistoni, G. Barsanti, P. F. Conte, and M. Del Tacca. Comparative pharmacokinetic analysis of 5-fluorouracil and its major metabolite 5-fluoro-5,6-dihydrouracil after conventional and reduced test dose in cancer patients. Clinical Cancer Research: An Official Journal of the American Association for Cancer Research, 6(8):3032–3037, August 2000. ISSN 1078-0432.

68. G. Codacci-Pisanelli, H.M. Pinedo, J. Lankelma, C.J. Van Groeningen, A.B.P. Van Kuilenburg, A.H. Van Gennip, and G.J. Peters. Pharmacokinetics of Bolus 5-Fluorouracil: Relationship Between Dose, Plasma Concentrations, Area-Under-the-Curve and Toxicity. Journal of Chemotherapy, 17(3):315–320, June 2005. ISSN 1120-009X, 1973-9478. doi: 10.1179/joc.2005.17.3.315.

69. A. Di Paolo, R. Danesi, A. Falcone, L. Cionini, F. Vannozzi, G. Masi, G. Allegrini, E. Mini, G. Bocci, P.F. Conte, and M. Del Tacca. Relationship between 5-fluorouracil disposition, toxicity and dihydropyrimidine dehydrogenase activity in cancer patients. Annals of Oncology, 12(9):1301–1306, September 2001. ISSN 09237534. doi: 10.1023/A:1012294617392.

70. Jennifer Saam, Gregory C. Critchfield, Stephanie A. Hamilton, Benjamin B. Roa, Richard J. Wenstrup, and Rajesh R. Kaldate. Body Surface Area–based Dosing of 5-Fluoruracil Results in Extensive Interindividual Variability in 5-Fluorouracil Exposure in Colorectal Cancer Patients on FOLFOX Regimens. Clinical Colorectal Cancer, 10(3):203–206, September 2011. ISSN 15330028. doi: 10.1016/j.clcc.2011.03.015.

71. Hideo Matsumoto, Hideo Okumura, Haruaki Murakami, Hisako Kubota, Masaharu Higashida, Atsushi Tsuruta, Kaoru Tohyama, and Toshihiro Hirai. Fluctuation in Plasma 5-Fluorouracil Concentration During Continuous 5-Fluorouracil Infusion for Colorectal Cancer. Anticancer Research, 35(11):6193–6199, November 2015. ISSN 1791-7530.

72. Andre Farkouh, Werner Scheithauer, Philipp Buchner, Apostolos Georgopoulos, Johannes Schueller, Birgit Gruenberger, and Martin Czejka. Clinical pharmacokinetics of capecitabine and its metabolites in combination with the monoclonal antibody bevacizumab. Anticancer Research, 34(7):3669–3673, July 2014. ISSN 1791-7530.

73. B. Reigner, K. Blesch, and E. Weidekamm. Clinical pharmacokinetics of capecitabine. Clinical Pharmacokinetics, 40(2):85–104, 2001. ISSN 0312-5963. doi: 10.2165/00003088-200140020-00002.

74. Yu Guo, Bing-Hong Xiong, Tao Zhang, Yong Cheng, and Li Ma. XELOX vs. FOLFOX in metastatic colorectal cancer: An updated meta-analysis. Cancer Investigation, 34(2):94–104, February 2016. ISSN 0735-7907, 1532-4192. doi: 10.3109/07357907.2015.1104689.

75. Michel Ducreux, Jaafar Bennouna, Mohamed Hebbar, Marc Ychou, Gérard Lledo, Thierry Conroy, Antoine Adenis, Roger Faroux, Christine Rebischung, Loic Bergougnoux, Leila Kockler, and Jean-Yves Douillard. Capecitabine plus oxaliplatin (XELOX) versus 5-fluorouracil/leucovorin plus oxaliplatin (FOLFOX-6) as first-line treatment for metastatic colorectal cancer. International Journal of Cancer, 128(3):682–690, February 2011. ISSN 0020-7136, 1097-0215. doi: 10.1002/ijc.25369.

76. FDA. FDA Oncology Center of Excellence: Project Optimus., 2021.

77. FDA. FDA. Optimizing the dosage of human prescription drugs and biological products for the treatment of oncologic diseases., 2024.

78. David S. Hong, Marwan G. Fakih, John H. Strickler, Jayesh Desai, Gregory A. Durm, Geoffrey I. Shapiro, Gerald S. Falchook, Timothy J. Price, Adrian Sacher, Crystal S. Denlinger, Yung-Jue Bang, Grace K. Dy, John C. Krauss, Yasutoshi Kuboki, James C. Kuo, Andrew L. Coveler, Keunchil Park, Tae Won Kim, Fabrice Barlesi, Pamela N. Munster, Suresh S. Ramalingam, Timothy F. Burns, Funda Meric-Bernstam, Haby Henary, Jude Ngang, Gataree Ngarmchamnanrith, June Kim, Brett E. Houk, Jude Canon, J. Russell Lipford, Gregory Friberg, Piro Lito, Ramaswamy Govindan, and Bob T. Li. KRAS ^g12c^ Inhibition with Sotorasib in Advanced Solid Tumors. New England Journal of Medicine, 383(13):1207–1217, September 2020. ISSN 0028-4793, 1533-4406. doi: 10.1056/NEJMoa1917239.

79. B.T. Hennessy, A.M. Gauthier, L.B. Michaud, G. Hortobagyi, and V. Valero. Lower dose capecitabine has a more favorable therapeutic index in metastatic breast cancer: retrospective analysis of patients treated at M. D. Anderson Cancer Center and a review of capecitabine toxicity in the literature. Annals of Oncology, 16(8):1289–1296, August 2005. ISSN 09237534. doi: 10.1093/annonc/mdi253.

80. Luke K. Fostvedt, Jennifer E. Hibma, Joanna C. Masters, Erik Vandendries, and Ana Ruiz-Garcia. Pharmacokinetic/Pharmacodynamic Modeling to Support the Re-approval of Gemtuzumab Ozogamicin. Clinical Pharmacology & Therapeutics, 106(5):1006–1017, November 2019. ISSN 0009-9236, 1532-6535. doi: 10.1002/cpt.1500.

81. Jack W. Scannell, James Bosley, John A. Hickman, Gerard R. Dawson, Hubert Truebel, Guilherme S. Ferreira, Duncan Richards, and J. Mark Treherne. Predictive validity in drug discovery: what it is, why it matters and how to improve it. Nature Reviews Drug Discovery, 21(12):915–931, December 2022. ISSN 1474-1776, 1474-1784. doi: 10.1038/s41573-022-00552-x.

82. Desamparados Roda, Begoña Jimenez, and Udai Banerji. Are Doses and Schedules of Small-Molecule Targeted Anticancer Drugs Recommended by Phase I Studies Realistic? Clinical Cancer Research, 22(9):2127–2132, May 2016. ISSN 1078-0432, 1557-3265. doi: 10.1158/1078-0432.CCR-15-1855.

83. Heinrich J. Huber and Hitesh B. Mistry. Explaining in-vitro to in-vivo efficacy correlations in oncology pre-clinical development via a semi-mechanistic mathematical model. Journal of Pharmacokinetics and Pharmacodynamics, 51:169–185, November 2024. ISSN 1567-567X, 1573-8744. doi: 10.1007/s10928-023-09891-7.

84. Pascal Schulthess, Vivi Rottschäfer, James W. T. Yates, and Piet H. Van Der Graaf. Optimization of Cancer Treatment in the Frequency Domain. The AAPS Journal, 21(6):106, November 2019. ISSN 1550-7416. doi: 10.1208/s12248-019-0372-4.

